# Variability in group size and the evolution of collective action

**DOI:** 10.1101/021485

**Authors:** Jorge Peña, Georg Nöldeke

## Abstract

Models of the evolution of collective action typically assume that interactions occur in groups of identical size. In contrast, social interactions between animals occur in groups of widely dispersed size. This article models collective action problems as two-strategy multiplayer games and studies the effect of variability in group size on the evolution of cooperative behavior under the replicator dynamics. The analysis identifies elementary conditions on the payoff structure of the game implying that the evolution of cooperative behavior is promoted or inhibited when the group size experienced by a focal player is more or less variable. Similar but more stringent conditions are applicable when the confounding effect of size-biased sampling, which causes the group-size distribution experienced by a focal player to differ from the statistical distribution of group sizes, is taken into account.

## 1 Introduction

Fish schools, wolf packs, bird flocks, and insect colonies exemplify the inherent tendency of animals to aggregate and live in groups (Krause and Ruxton, 2002; Sumpter, 2010). Within these groups, animals engage in a vast array of collective actions such as foraging (Giraldeau and Caraco, 2000), hunting (Packer and Ruttan, 1988), vigilance (Ward et al., 2011), defense (Hartbauer, 2010), and navigation (Simons, 2004). These social interactions are not without conflict, as individual and collective interests can oppose each other to the point of discouraging joint action and the pursuit of common goals.

Here we follow the game-theoretic approach of modelling such social dilemmas involved in collective action as multiplayer matrix games in which payoffs for individuals are determined by their own action, namely whether to cooperate or not, and the number of other individuals within their group who choose to cooperate (Broom et al., 1997; Peña et al., 2014). As shown in the vast literature on nonlinear public goods games (e.g., Dugatkin, 1990; Motro, 1991; Bach et al., 2006; Hauert et al., 2006; Cuesta et al., 2008; Pacheco et al., 2009; Archetti and Scheuring, 2011) cooperative behavior may arise in the evolutionary solution of such games even when other mechanisms potentially promoting cooperation such as relatedness (Eshel and Motro, 1988; Archetti, 2009; Peña et al., 2015) and reciprocity in repeated interactions (Boyd and Richerson, 1988; Hilbe et al., 2014) are absent.

Evolutionary models of collective action, including the ones cited above, typically assume that social interactions occur in groups of identical size. In contrast, empirical studies of animal group sizes show large variation in group size (Bonabeau et al., 1999; Gerard et al., 2002; Jovani and Tella, 2007; Griesser et al., 2011; Hayakawa and Furuhashi, 2012). This paper studies how this intrinsic variability in group size affects the evolution of cooperative behavior. We do so by modeling the evolutionary dynamics with the replicator dynamics (Taylor and Jonker, 1978; Hofbauer and Sigmund, 1998) and under the assumptions that the group-size distribution is exogenous, the population is well-mixed, and individuals express one of the two possible pure strategies. This is the same setting as the one used in Peña (2012) to investigate the effects of group-size diversity in public goods, that is, without any frequency-dependent or assortment bias in group composition. Although real group formation processes will certainly lead to such biases, we stick to this setting as it allows us to infer the consequences of relaxing the assumption of fixed group sizes without introducing the confounding effect of strategy assortment.

We identify general conditions, both on the class of group-size distributions and on the payoff structure of the collective action problem, which allow us to conclude whether more or less variation in group size promotes or inhibits cooperation. We thus go beyond Peña (2012) in not limiting us to the comparison of a deterministic group size with a variable group size (resp. the comparison of three particular group-size distributions) and by going beyond particular examples for collective action problems such as the volunteer’s dilemma (Diekmann, 1985) and public goods game with synergy or discounting (Hauert et al., 2006).

To obtain our results, we combine three different kinds of insights. First, we build on results obtained in Motro (1991) and Peña et al. (2014) to identify conditions on the payoff structure of the game which are sufficient to infer those shape properties of the gain function that are required to identify the variability effects we are interested in (Lemmas 1 and 2). These results dispense with the need to explicitly calculate the gain function (i.e., the difference in expected payoff between the two strategies) whenever the payoff structure of the game satisfies the relevant conditions.

Second, we use the theory of stochastic orders (Shaked and Shanthikumar, 2007) to give precise meaning to the notion that one distribution is more ore less variable than another. This allows us to extend the comparison between a deterministic group size and a variable group size considered in Peña (2012) to the comparison of different group-size distributions. In particular, the very same condition on the shape of the gain function (when viewed as a function of group size) which Peña (2012) identified as being sufficient for group-size variability to promote cooperation relative to the benchmark of a deterministic group size yields the same conclusion for any two group-size distributions that can be compared in the convex order (Shaked and Shanthikumar, 2007). Many commonly considered group-size distributions with the same expected value can be compared in this way and often this is easy to check graphically.

Third, we demonstrate that focusing on the variability of the group-size distribution *per se* confounds two effects that are better understood when viewed separately. The issue is that the proportion of groups with a given size *s* is not identical to the proportion of individuals in groups with size *s* because a randomly chosen individual is more likely to find itself in a large rather than a small group. Whereas the former proportions are described by the group-size distribution, the latter are described by the so-called size-biased sampling distribution (Patil and Rao, 1978) that, for convenience, we refer to as the experienced group-size distribution. The empirical importance of distinguishing the group-size distribution and the experienced group-size distribution is well-understood in the statistical literature; a recent discussion in a biological context can be found in Jovani and Mavor (2011). The theoretical importance of distinguishing between the two distributions in our setting arises because an increase in the variability of the experienced group-size distribution may have different evolutionary consequences than an increase in the variability of the group-size distribution. This is because more variability in group size does not simply induces more variability in experienced group size but also increases average experienced group size.

Our main results are summarized in Propositions 1 and 2. These propositions are stated in terms of the gain sequence of the game, which collects the gains from switching (Peña et al., 2014), i.e., the difference in payoff a focal player obtains from switching its strategy as a function of the number of other cooperating players in the focal player’s group. Proposition 1 shows that more variation in experienced group size promotes the evolution of cooperative behavior whenever the payoff structure of the game is such that the gain sequence is convex, whereas with concave gains from switching more variation in experienced group size inhibits the evolution of cooperative behavior. ^1^ Because more variation in group size not only implies more variation in experienced group size but also an upward shift in the experienced group-size distribution, these conditions do not suffice to imply that more variation in group size (rather than in experienced group-size) promotes or inhibits cooperative behavior. Proposition 2 takes this confounding effect into account and shows that more variation in group size promotes cooperative behavior whenever the gain sequence is convex and increasing, whereas cooperative behavior is inhibited when the gain sequence is concave and decreasing.

The difference between the sufficient conditions in Propositions 1 and 2 is significant as there are interesting collective action problems for which the gains from switching are convex or concave but fail the additional monotonicity properties required to determine whether more variation in group size promotes or inhibits cooperation. We illustrate this and other features of our analysis by using the volunteer’s dilemma (Diekmann, 1985) and the public goods game with synergy or discounting (Hauert et al., 2006, Section 2.3.2) as examples. Further examples will be provided in Section 4, where we also discuss classes of collective action problems for which our approach is not applicable because the gain sequences are neither convex nor concave. Finally, we investigate the consequences of our main results for the number and location of stable rest points of the replicator dynamics, demonstrating that an increase or decrease in experienced group-size variability can induce transcritical and saddle-node bifurcations by which rest points can be created, destroyed, and their stability changed.

## 2 Methods

### 2.1 Group size and experienced group size

We consider an infinitely large and well-mixed population subdivided into groups consisting of a finite number of individuals. We assume that group size is given by a random variable *S* with support in the non-negative integers, probability distribution *p* = (*p*_0_, *p*_1_, . . .), and finite expected value 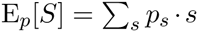. We refer to *p* as the group-size distribution and assume throughout that *p*_0_ + *p*_1_ < 1 holds, so that the fraction of groups with at least two individuals is not zero.^2^

Given a group-size distribution *p*, the fraction 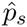 of individuals who find themselves in a group of size *s* ≥ 1 is

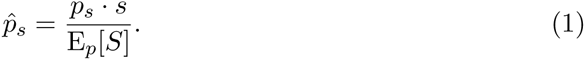

We refer to the probability distribution 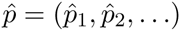 defined by (1) as the experienced group-size distribution and to its associated random variable *Ŝ* as the experienced group size. In the statistical literature the experienced group-size distribution is known as the size-biased sampling distribution (Patil and Rao, 1978).

Unless group size is deterministic, the experienced group-size distribution differs from the group-size distribution because a randomly sampled individual is more likely to be a member of a large group than of a small group. Table 1 shows the relationship between group size and experienced group size for some distributions that are commonly used to model variation in group size, including the classical models of Poisson and negative binomial distributions (Okubo, 1986) and the logarithmic distribution featured in recent theoretical and empirical work on animal group-size distributions (Niwa, 2003; Ma et al., 2011; Griesser et al., 2011). We will employ the distributions from Table 1 to illustrate our subsequent analysis.

**Table 1:** Experienced group-size distributions (*Ŝ*) for some common group-size distributions (*S*). Po(λ) refers to a Poisson distribution with parameter λ > 0, which has support on the nonnegative integers and expected value λ. ΝΒ(*η*, *π*) refers to a negative binomial distribution with parameters *η* > 0 and 0 < *π* < 1, which has support on the nonnegative integers and expected value *ηπ*/(1 – *π*). L(*δ*) refers to a logarithmic distribution with parameter 0 < *δ* < 1, which has support on the natural numbers and expected value *δ*/ [(*δ* – 1) ln(1 – *δ*)]. Note that ΝΒ(1, *π*) corresponds to a geometric distribution. See Table 1 in Patil and Rao (1978) for these and further examples.

| $S$ | $p_s$ | $\hat{S}$ | $\hat{p}_s$ for $s \geq 1$ |
| --- | --- | --- | --- |
| Po($\lambda$) | $\frac{\lambda^s}{s!} \exp(-\lambda)$ | $1 + \text{Po}(\lambda)$ | $\frac{\lambda^{s-1}}{(s-1)!} \exp(-\lambda)$ |
| NB($\eta, \pi$) | $\binom{\eta+s-1}{s} \pi^s (1-\pi)^\eta$ | $1 + \text{NB}(\eta+1, \pi)$ | $\binom{\eta+s-1}{s-1} \pi^{s-1} (1-\pi)^{\eta+1}$ |
| L($\delta$) | $\frac{-1}{\ln(1-\delta)} \frac{\delta^s}{s}$ | $1 + \text{NB}(1, \delta)$ | $\delta^{s-1} (1-\delta)$ |

### 2.2 Social interactions and gain sequence

Social interactions take place within groups: individuals within each group of size *s* ≥ 1 participate in a symmetric *s*-player game. In this game individuals either cooperate (play action *A*, contribute to the provision of a collective good) or defect (action *B*, do not contribute to the provision of a collective good). The payoff for an individual is determined by its own action and the number of other individuals in the group who play action A. Let *a_k_* denote the payoff to an A-player and *b_k_* denote the payoff to a *B*-player when *k* = 0, 1,..., *s* – 1 co-players play *A* (and hence *s* – 1 – *k* co-players play B). We alternatively refer to *A*-players as “cooperators” and *B*-players as “defectors”.

Let *d_k_* = *a_k_* – *b_k_* denote the *k*-th gain from switching, i.e., the gain in payoff an individual makes from cooperating rather than defecting when *k* co-players cooperate, and let *d* = (*d*_0_, *d*_1_,...) denote the corresponding gain sequence. The gain sequence *d* is increasing (decreasing, convex, concave) if Δ*d_k_* ≥ 0 (Δ*d_k_* ≤ 0, Δ^2^*d_k_* ≥ 0, Δ^2^*d_k_* ≤ 0) holds for all *k* ≥ 0, where Δ*d_k_* = *d_k_*_+1_ – *d_k_* and Δ^2^*d_k_* = Δ*d_k_*_+1_ – Δ*d_k_*. Examples 1 and 2 below, based on Diekmann (1985) and Hauert et al. (2006, Section 2.3.2), illustrate how these properties of gain sequences arise in two familiar collective action games. Peña et al. (2014) provide further examples and general discussion of gain sequences, their properties, and their importance for the evolutionary analysis of multiplayer games.

#### Example 1

(Volunteer’s dilemma). In the volunteer’s dilemma each cooperator pays a cost *c* > 0, whereas defectors incur no cost. If there is at least one cooperator (“volunteer”) in the group, a public good is produced that provides a benefit *u* > *c* to each member of the group. If there are no cooperators in the group, payoffs are zero for all individuals in the group. The payoffs in this game are given by *a_k_* = (*u* – *c*, *u* – *c*, *u* – *c*,...) and *b_k_* = (0, *u*, *u*,...). The gain sequence is thus *d_k_* = (*u* – *c*, –*c*, –*c*,...). Here Δ*d_k_* = (–*u*, 0, 0,...) and Δ^2^*d_k_* = (*u*, 0, 0,...), so that the gain sequence is decreasing and convex.

#### Example 2

(Public goods game with synergy or discounting). As in the volunteer’s dilemma each cooperator incurs a cost *c* > 0 for a public good to be produced. In contrast to the volunteer’s dilemma, the benefit each group member obtains from the public good depends on the number of cooperators in the group and may also differ between cooperators and defectors. Specifically, if there are *j* ≥ 1 cooperators in the group, the value of the public good is 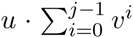 for defectors and 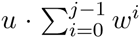 for cooperators, where *u* > 0, *v* > 0, and *w* > 0 are parameters. The gain sequence for this social interaction is ^3^

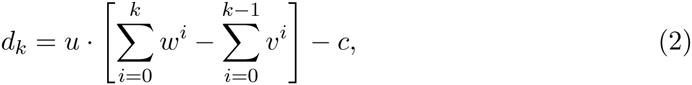

so that we have

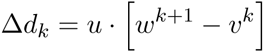

and

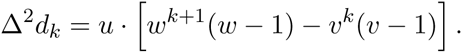

If *w* = *v* holds (that is, cooperators and defectors obtain the same benefit), the gain sequence is increasing and convex for *w* = *v* > 1 and is decreasing and convex for *w* = *v* < 1. More generally, the gain sequence is increasing and convex if *w* ≥ 1 and *w* ≥ *v* holds and is decreasing and convex if 1 ≥ *v* ≥ *w* and *w*(1 – *w*) ≤ 1 – *v* holds. For other parameter values the gain sequence may have different shapes. In particular, if either *v* ≥ 1 ≥ *w* holds or the conditions 1 ≥ *w* ≥ *v* and *w*(1 – *w*) ≥ 1 – *v* are both satisfied, the gain sequence is decreasing and concave. If *v* ≥ *w* ≥ 1 and *w*(*w* – 1) ≤ *v* – 1 holds, the gain sequence is concave and unimodal (that is, increasing up to some critical value of *k* and decreasing thereafter).

Before proceeding, we note that the game introduced in Example 2 differs from the one introduced in Hauert et al. (2006) —and studied in Peña (2012) for the case *v* = *w*— in that the benefits obtained from the public good are not scaled by the inverse of the group size. We return to this point in Section 4.

### 2.3 Gain function and expected gain function

If the proportion of *A*-players in the population is *x*, the average payoffs obtained by an *A*-player and a *B*-player who find themselves in a group of size *s* are respectively given by

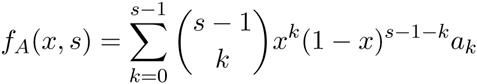

and

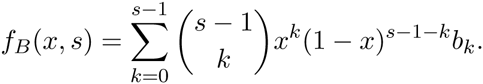

The difference between the average payoff of *A*-players and *B*-players in groups of size *s* is then

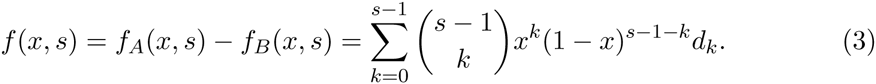

The difference between the average payoff of an *A*-player and a *B*-player in the population is the expectation of *f* (*x*, *Ŝ*) and thus given by

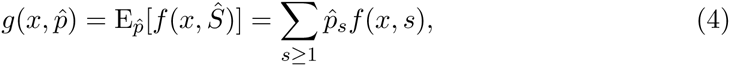

where we use the subscript 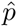 on the expectation operator to emphasize its dependance on the experienced-group size distribution. Throughout the following we refer to *f* (*x*, *s*) as the gain function and to 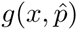 as the expected gain function.

Defining

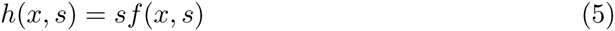

and using (1) the expected gain can be rewritten in terms of the underlying group-size distribution as

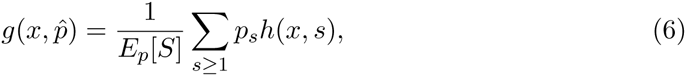

which is the expression used by Peña (2012, Eq. 3).

### 2.4 Evolutionary dynamics

We assume that the change in frequency of A-players in the population is described by the replicator dynamics (Taylor and Jonker, 1978; Hofbauer and Sigmund, 1998)

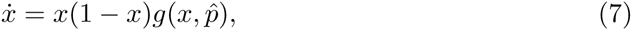

while noting that any other dynamics in which the direction of selection (i.e., the sign of *ẋ*) is determined by the sign of the expected gain function in the same way as for the replicator dynamics will lead to identical results.

The replicator dynamics has two rest points at *x* = 0 (where the whole population consists of defectors) and at *x* = 1 (where the whole population consists of cooperators). Interior rest points are given by the values *x*^*^ ∈ (0, 1) satisfying 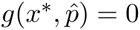. An interior rest point *x*^*^ is stable if the expected gain function changes its sign from positive to negative at *x*^*^, for which 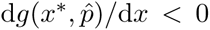 is a sufficient condition. Regarding the endpoints, *x* = 1 is stable if the expected gain is positive for sufficiently large *x*, for which 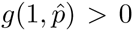 is a sufficient condition. Similarly, *x* = 0 is stable if the expected gain is negative for sufficiently small *x*, for which 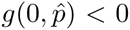 is a sufficient condition. Because *f* (0, *s*) = *d*_0_ holds for all *s* we have

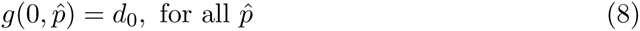

so that the stability of the rest point *x* = 0 does not depend on the group-size distribution. To simplify the exposition, we assume *d*_0_ ≠ 0 throughout the following.

When group-size is deterministic and given by *s*, (7) reduces to

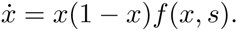

This is the version of the replicator dynamics considered in Peña et al. (2014), who show how shape properties of the gain sequence *d* can be used to infer shape properties of the gain function *f* and, thus, information about the number and stability of the rest points of the replicator dynamics for a given deterministic group size.

To illustrate the relationship between the gain sequence, the (expected) gain function, and the rest points of the replicator dynamics, let us consider the volunteer’s dilemma from Example 1. Substituting the gain sequence *d* = (*u* - *c*, -*c*, -*c*,...) into (3) yields the gain function *f* (*x*, *s*) = *u*(1 - *x*)*^s^*^-1^ - *c*. The gain function is strictly decreasing in *x* and satisfies *f* (0, *s*) > 0 as well as *f* (1, *s*) < 0, so that there is exactly one interior rest point *x*^*^, which is also the unique stable rest point of the replicator dynamics when all groups have identical size *s*. The expected gain function, given by 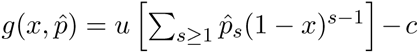, is also strictly decreasing in *x*. Further, 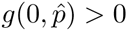 holds and, provided that 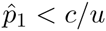 holds (meaning that an individual is not too likely to find itself in the position of being the sole member of a group), we also have 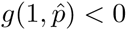. Hence, when the experienced group-size distribution is 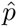, the replicator dynamics will again have one interior rest point *x*^*^, which is also the unique stable rest point of the dynamics. While for deterministic group sizes this stable rest point is easily calculated as *x*^*^ = 1 - (*c/u*)^1/(^*^s^*^-1)^ (Diekmann, 1985), even for a game as simple as the volunteer’s dilemma no analytical solution for the stable rest point can be determined for general group-size distributions. Nevertheless, once the right tools are brought to bear on the issue, a great deal can be said not only about the impact of variability in experienced group size on the evolutionary dynamics for the volunteer’s dilemma but also for more complicated games such as the one considered in Example 2.

### 2.5 Variability order

As our interest is in isolating the effect of variation in group size on the evolutionary dynamics, we have to take a stance on how to compare the variability of two distributions. We follow the standard approach from the literature on stochastic orders (Shaked and Shanthikumar, 2007) and consider one (experienced) group-size distribution to be more variable than another if it is more “spread out” in the sense of the so-called convex order. Throughout the following we write *q* ≥*_v_ p* if group-size distribution *q* is more variable than *p* in this sense and similarly write 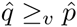 if the experienced group-size distribution 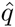 associated with *q* is more variable than the experienced group-size distribution 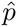 associated with *p*.

By definition (Shaked and Shanthikumar, 2007, Chapter 3), *q* ≥*_v_ p* means that for all convex functions *φ*: ℝ → ℝ the inequality E*_q_*[*φ*(*Υ*)] ≥ E*_p_*[*φ*(*Χ*)] holds. As *φ*(*x*) = *x* and *φ*(*x*) = - *x* are both convex functions, *q* ≥*_v_ p* implies E*_q_*[*Y*] = E*_p_*[*X*]. As *φ*(*x*) = *x*^2^ is a convex function, *q* ≥*_v_ p* implies Var*_q_*[*Y*] ≥ Var*_p_*[*X*]. Consequently, a necessary condition for a distribution *q* to be more variable than a distribution *p* is that *q* and *p* have the same expected value and that the variance of *q* is at least as high as the variance of *p*. Of course (here and in the following discussion of sufficient conditions) the same statements are applicable for experienced group-size distributions.

The conditions E*_q_*[*Y*] = E*_p_* [*X*] and Var*_q_* [*Y*] ≥ Var*_p_*[*X*] are not sufficient to imply that *q* is more variable in the convex order than *p*. Rather, provided that the expected values are the same, a sufficient condition for *q* ≥*_v_ p* is that *q* assigns higher probability to more extreme realizations of group size in the sense that the sequences (*p*_0_, *p*_1_,...) and (*q*_0_, *q*_1_,...) cross exactly twice with *q_s_* > *p_s_* holding for *s* sufficiently small and *s* sufficiently large, whereas *p_s_* > *q_s_* holds for intermediate values of *s* (Shaked and Shanthikumar, 2007, p. 133). This sufficient condition is trivially satisfied when *p* describes a deterministic group size: any group-size distribution *q* with expected value *s* is more variable than the deterministic group-size distribution *p* assigning probability 1 to *s*. Less trivially, all the (experienced) group-size distributions appearing in Table 1 are ordered by variability when their expected values coincide: As we show in Appendix A.1, negative binomial distributions with the same expected value are ordered by variability according to the value of the parameter *η*, with the geometric distribution (corresponding to the case *η* = 1) being most variable and the Poisson distribution (corresponding to the limit case *η* → ∞) being least variable. See Fig. 1 for an illustration. The logarithmic distribution is even more variable than the geometric distribution. The truncated Poisson, geometric, and Waring distributions considered in Peña (2012, Fig. 1) provide further examples of distributions satisfying the sufficient condition stated above. It follows that all the comparisons considered in Peña (2012) are ones in which the group-size distributions are ordered by variability.

**Figure 1:**
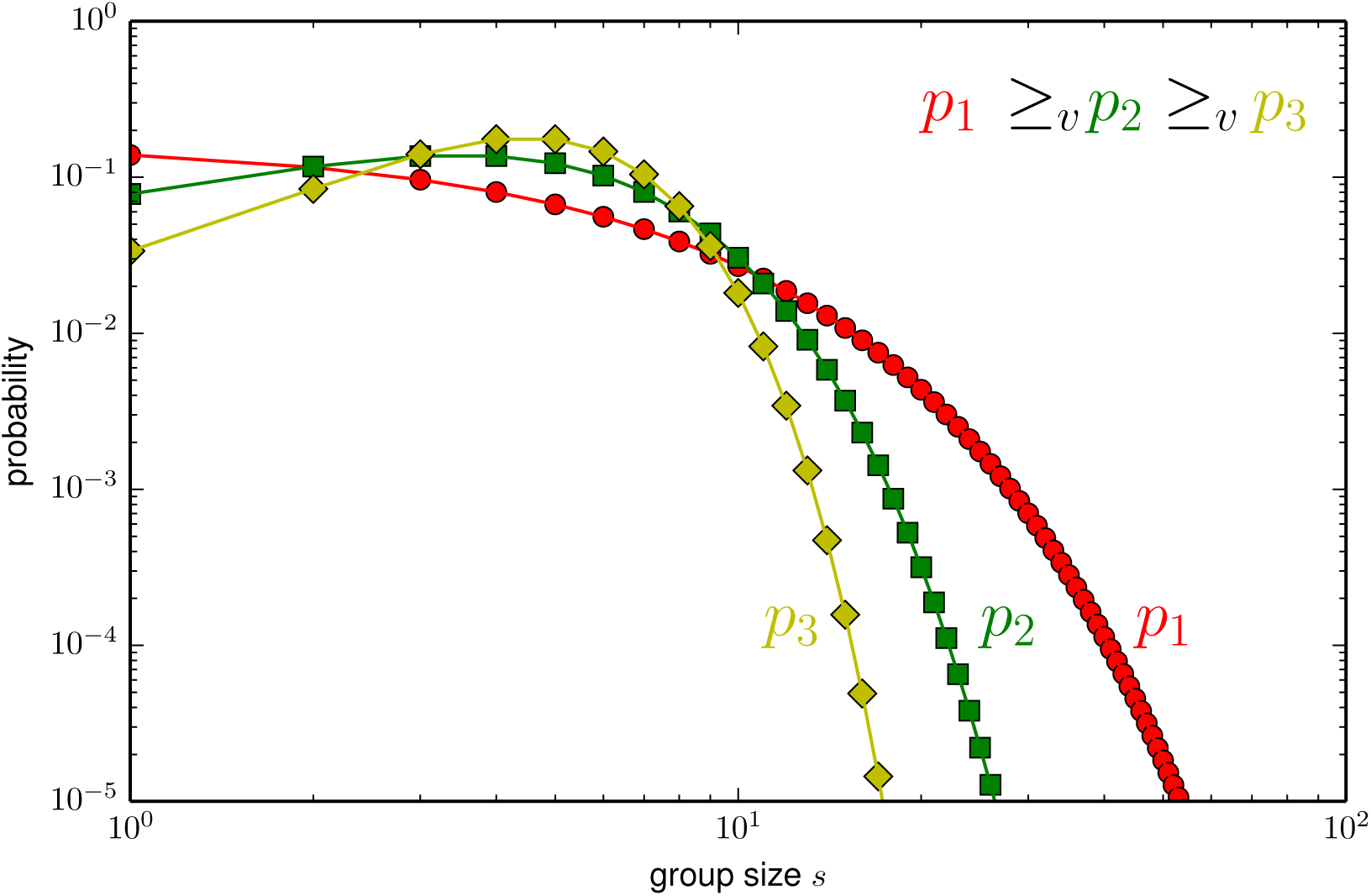
Three group-size distributions ordered by variability. *p*_1_ is the geometric distribution NB(1, 5/6), *p*_2_ the negative-binomial distribution NB(5, 1/2), and *p*_3_ is the Poisson distribution Po(5). All three distributions have an expected value of 5 and are thus, as shown in Appendix A.1, ordered by variability with the geometric distribution *p*_1_ being most variable and the Poisson distribution *p*_3_ being least variable. As explained in the text this can be seen graphically by observing that each pair of probability mass functions crosses exactly twice, with the geometric distribution assigning most weight and the Poisson distribution assigning least weight to extreme realizations.

One might think of pursuing the simpler approach of considering one of two distributions with the same expected value to be “more variable” than the other if it has the higher variance. Alas, such an approach would be of very limited applicability in our context: unless the expected gain function is determined by the first two moments of the experienced group-size distribution, knowledge of the expected value and variance of the experienced group-size distribution (or the actual group-size distribution for that matter) does not provide enough information to determine the expected gain function.^4^ This precludes the possibility of obtaining any general results linking the variance of the experienced group size to the evolutionary dynamics. Fig. 2 illustrates this for the volunteer’s dilemma.

**Figure 2:**
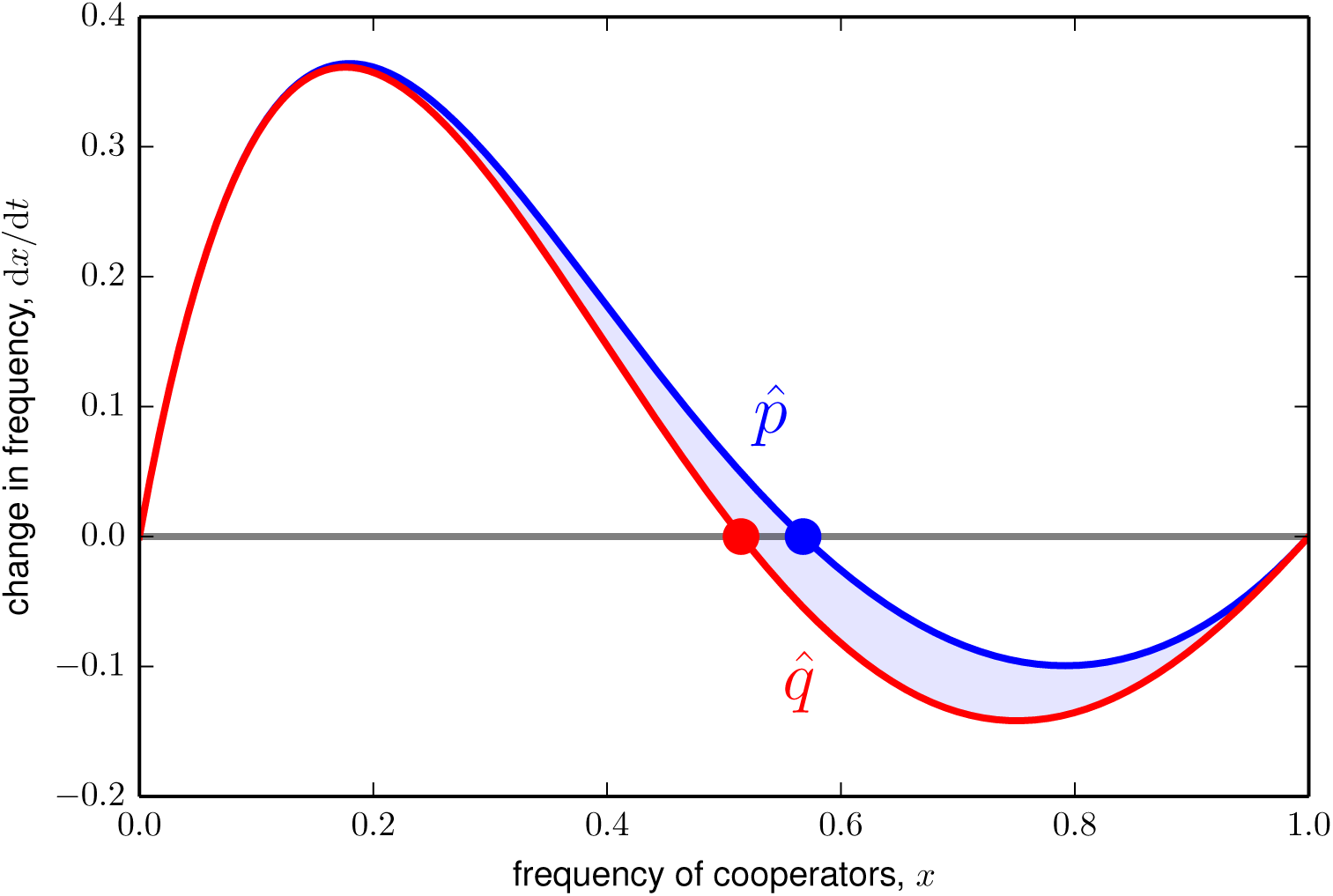
Different experienced group-size distributions with the same mean and variance can lead to different evolutionary dynamics. Here we illustrate the evolutionary dynamics as given by (7) for the volunteer’s dilemma with *c* = 1, *u* = 6 (cf. Example 1) and two experienced group-size distributions. The first distribution, 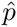 (blue), has support {2, 4, 6} with 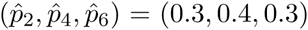; the second distribution, 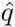 (red), has support {3, 4, 7} with 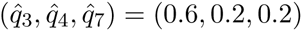. With these values, 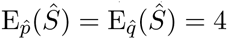 and 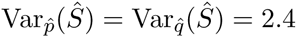. The replicator dynamics for these two cases are however different, with the distribution 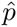 leading to the stable rest point 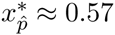 (*blue circle*), and the distribution 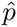 leading to the stable rest point 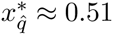 (*red circle*).

### 2.6 Variability effects and experienced variability effects

A final complication we must face before proceeding to our results is that the distinction between a more variable group size and a more variable experienced group size is nontrivial. The reason is that the two conditions E*_p_*[*S*] = E*_q_* [*S*] and 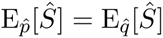 are not equivalent. Indeed, a simple calculation shows that for any group-size distribution *p* we have

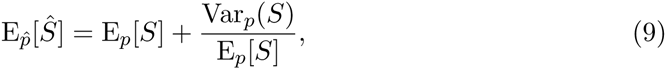

where Var*_p_*(*S*) denotes the variance of the group-size distribution. Hence, whenever two group-size distributions *p* and *q* satisfy *q* ≥*_v_ p* and *q* has a strictly higher variance than *p*, the experienced group-size distribution 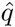 has a strictly higher expected value than the experienced group-size distribution 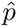. Consequently, *q* ≥*_v_ p* does not imply 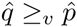, as the two experienced group-size distributions will not have the same expected value.

Fig. 3 illustrates the importance of this point for the case of the volunteer’s dilemma and negative-binomially distributed group-sizes. The left panel of the figure shows that the frequency of cooperators at the stable rest point is a decreasing function of group-size variability, whereas the right panel shows that the frequency of cooperators at the stable rest point is an increasing function of experienced group-size variability. As will become clear later, these strikingly different effects of a change in variability are entirely driven by the higher average experienced group sizes associated with more variable group sizes.

**Figure 3:**
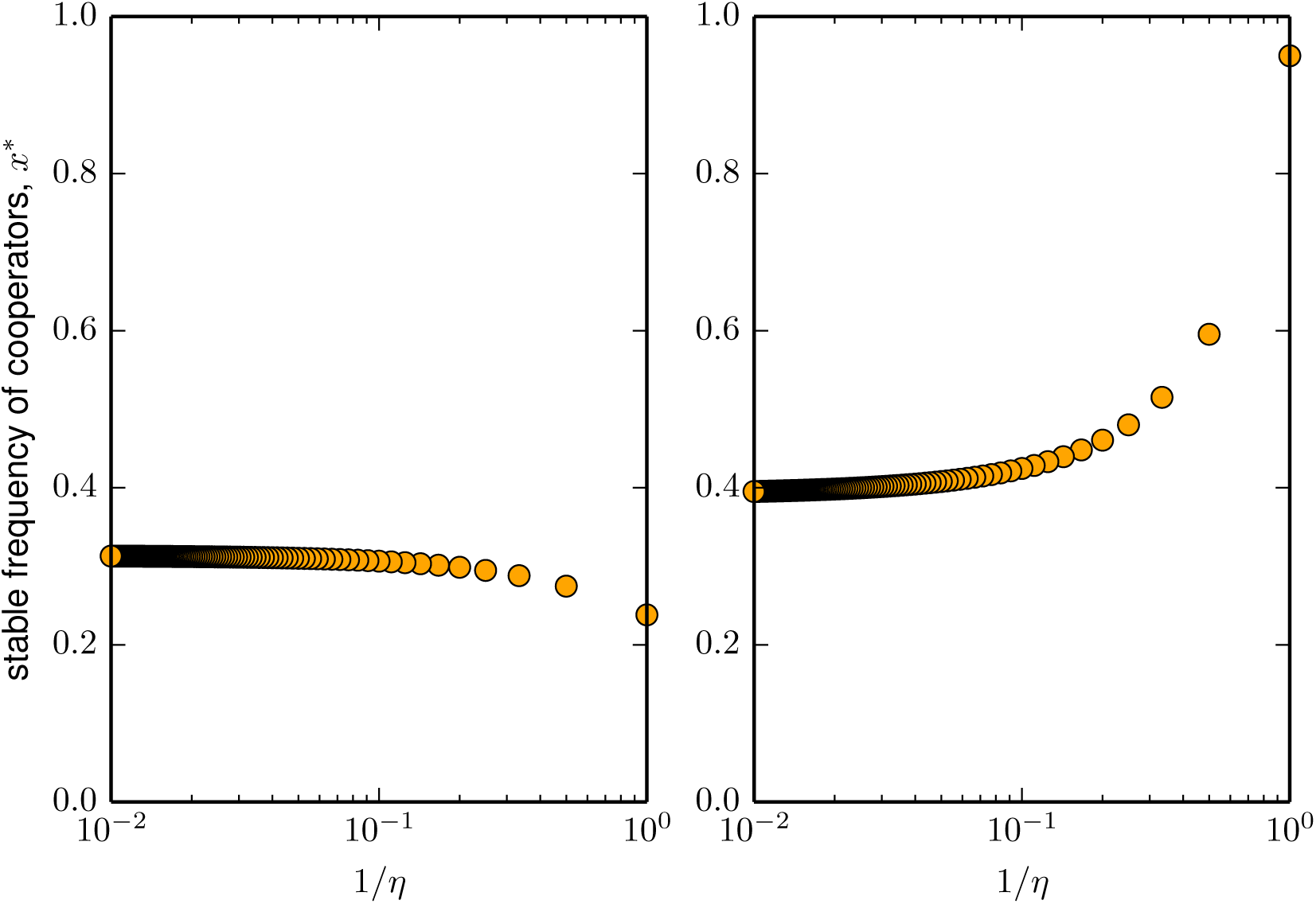
Stable frequency of cooperators *x*^*^ as a function of group-size variability (*left*) and experienced group-size variability (*right*) in the volunteer’s dilemma (*c* = 2.5, *u* = 12). In both panels variability increases when going from left to right with the inverse of the parameter *η* of a negative-binomial group-size distribution on the horizontal axis (cf. Table 1). *Left:* group size is distributed according to the negative binomial ΝΒ(*η*, *π*) with parameter *π* adjusted such that the expected group size is 5 for all *η*. The stable fraction of cooperators is a decreasing function of group-size variability as measured by 1/*η*. *Right:* group size is distributed according to the negative binomial ΝΒ(*η*, *π*) with parameter *π* adjusted such that the expected experienced group size is 5 for all *η*. The stable fraction of cooperators is an increasing function of experienced group-size variability as measured by 1/*η*.

We respond by distinguishing between variability and experienced-variability effects on the frequency of *A*-players in the population. In particular, we say that there is a positive experienced-variability effect if 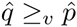 implies

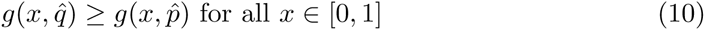

and a negative experienced-variability effect if 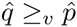 implies

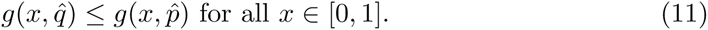

Hence, in the case of a positive (resp. negative) experienced-variability effect, more variability in experienced group size unambiguously increases (resp. decreases) the difference between the average fitness of cooperators and defectors. Similarly, we say that the variability effect is positive if *q* ≥*_v_ p* implies (10) and that it is negative if *q* ≥*_v_ p* implies (11). In either case, the interpretation is that (10) means that variability promotes the evolution of cooperation, whereas (11) means that variability inhibits the evolution of cooperation.

## 3 Results

Our analysis proceeds in four steps. First, we establish two preliminary results that relate shape properties of the gain sequence *d* to corresponding properties of the gain function *f* (*x*, *s*). Second, we identify conditions on the gain sequence *d* which allow us to sign the experienced-variability effect. Third, we turn to the more challenging task of signing the variability effect. Fourth, we draw out the implications of the inequalities in (10) and (11) for the number and location of the rest points of the replicator dynamics under the conditions which allow us to sign the experienced-variability effect.

### 3.1 Preliminaries

As noted and discussed in Peña et al. (2014), the gain function *f* (*x*, *s*) is a polynomial in Bernstein form. The following two preliminary results summarize the properties of the gain function and the expected gain function that are implied by the theory of polynomials in Bernstein form (Farouki, 2012) and are of relevance for our analysis. We begin by relating the monotonicity and convexity properties of the gain sequence

We begin by relating the monotonicity and convexity properties of the gain sequence *d* to corresponding properties of the gain function *f* (*x*, *s*) when considered as a function of group size *s*. Formally, we say that *f* (*x*, *s*) is increasing (resp. decreasing) in *s* if Δ*_s_f* (*x*, *s*) = *f* (*x*, *s* + 1) - *f* (*x*, *s*) ≥ 0 (resp. Δ*_s_f* (*x*, *s*) ≤ 0) holds for all *s* ≥ 1 and *x* ∈ [0,1]; *f* (*x*, *s*) is convex (resp. concave) in *s* if 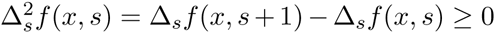 (resp. 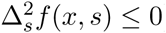) holds for all *s* ≥ 1 and *x* ∈ [0,1]. With this terminology in place, we can state the following lemma. The proof, which uses an observation due to Motro (1991), is in Appendix A.2.

#### Lemma 1.

*If the gain sequence d is increasing (decreasing, convex, concave), then the gain function f* (*x, s*) *is increasing (decreasing, convex, concave) in group size s.*

As noted in Peña et al. (2014, Remark 3), the gain function *f* (*x*, *s*) inherits the monotonicity and convexity properties of the gain sequence *d* when considered as a function of *x*. In particular, when the gain sequence *d* is increasing (decreasing), then the gain function *f* (*x*, *s*) is increasing (decreasing) in *x*. Similarly, when the gain sequence *d* is convex (concave), then *f* (*x*, *s*) is convex (concave) in *x*. As monotonicity and convexity properties are preserved by taking weighted averages, it is immediate from (4) that the expected gain function 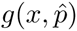 inherits these monotonicity and convexity properties in *x* no matter what the experienced group-size distribution 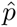 is. The following result thus requires no further proof.

#### Lemma 2.

*If the gain sequence d is increasing (decreasing, convex, concave), then the expected gain function* 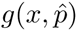 *is increasing (decreasing, convex, concave) in the proportion x of A-players for all experienced group-size distributions* 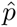.

### 3.2 Signing the experienced-variability effect

Suppose that the gain sequence *d* is convex. Then, as established in Lemma 1, the function *f* (*x*, *s*) is convex in group size *s* no matter what the fraction *x* of *A*-players in the population is. By the very definition of the relationship 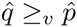, convexity of *f* (*x*, *s*) in group-size *s* is in turn sufficient to imply the inequality 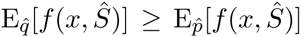 or, recalling the definition of the expected gain function in (4), the inequality in (10). It thus follows that the experienced-variability effect is positive whenever the gain sequence *d* is convex. An analogous argument shows that concavity of *d* is sufficient to imply inequality (11). Consequently, we obtain the following simple sufficient conditions on the payoff structure of the game under which the experienced-variability effect can be signed. The formal proof is in Appendix A.2.

#### Proposition 1

*[1.1] If the gain sequence d is convex, then the experienced-variability effect is positive.*

*[1.2] If the gain sequence d is concave, then the experienced-variability effect is negative.*

As we have noted before, the gain sequence for the volunteer’s dilemma in Example 1 is convex. Hence, it is an immediate implication of Proposition 1.1 that the experienced-variability effect is positive for the volunteer’s dilemma. The results for the public goods game with synergy or discounting in Example 2 are more nuanced here the gain sequence is convex for some parameter values (including the case *v* = *w* considered in Peña (2012)) and concave for others. From Proposition 1 an increase in experienced variability promotes cooperation in the former case but inhibits it in the latter.

### 3.3 Signing the variability effect

Heuristically, we may think of an increase in variability in group-size as giving rise to two effects, namely (i) an increase in the variability of the experienced group-size distribution and (ii) an upward shift in that distribution. As we have discussed in Section 2.6, the source of the second effect is that an increase in the variability of the group-size distribution increases the expected value of the experienced group size: *q* ≥*_v_ p* implies 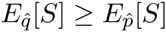.

Provided that the gain sequence is convex or concave, the first of these effects —the experienced variability effect— can be signed (Proposition 1). It is intuitive that the sign of the second effect, namely the size effect resulting from an increase in the expected experienced group size, is determined by the monotonicity properties of the gain sequence: for a given proportion *x* of *A*-players in the population a higher experienced group size increases the average number of *A*-players in a group a focal player will experience, which in turn increases (resp. decreases) the difference in average payoff between *A*-players and *B*-players when the gain sequence is increasing (resp. decreasing). This suggests that the variability effect can be signed if the gain sequence is either increasing and convex or decreasing and concave because in these cases the experienced-variability effect and the size effect both point in the same direction.

The proof of the following proposition in Appendix A.2 confirms this intuition. In this proof the function *h*(*x*, *s*) = *sf* (*x*, *s*), that we have introduced in (5), plays a central role. Writing the expected gain function as in (6) it is immediate from the definition of the convex order that convexity (resp. concavity) of the function *h*(*x*, *s*) is sufficient to imply a positive (resp. negative) variability effect. This first step of the proof generalizes the observation from Peña (2012) that convexity of *h*(*x*, *s*) implies that group-size variability promotes cooperation relative to the benchmark of a deterministic group size. We complete the proof by showing that the function *h*(*x*, *s*) is convex (resp. concave) in group size *s* for all *x* when the gain sequence is increasing and convex (resp. decreasing and concave).

#### Proposition 2

*[2.1] If the gain sequence d is increasing and convex, then the variability effect is positive.*

*[2.2] If the gain sequence d is decreasing and concave, then the variability effect is negative.*

While there are collective action problems for which the gain sequence satisfies the conditions appearing in Proposition 2 —for instance, Example 2 provides conditions on the parameter values of the public goods game with synergy or discounting for which this is the case— these conditions are much more stringent that the ones in Proposition 1. In the cases not covered by Proposition 2, e.g., when the gain sequence is decreasing and convex (as in the volunteer’s dilemma from Example 1) or is concave and unimodal (as it is the case in the model of Bach et al. (2006) or for a broad range of parameter values in the game considered in Example 2), no clear-cut prediction for the variability effect is possible. The reason is that in such games the experienced-variability effect and the size effect may not only point in opposite directions but their relative strength depends on the frequency *x* of cooperators in the population and, further, on the particular group-size distributions under consideration. As a consequence, even in the simplest case in which the replicator dynamics has a unique stable rest point, no general conclusions about the effect of an increase in group-size variability on the the location of this rest point are possible. For instance, while Fig. 3 documents a case in which the stable rest point in the volunteer’s dilemma is decreasing in group-size variability, it is apparent from Peña (2012, Fig. 5) that for other parameter values increasing group-size variability can either increase or decrease the stable frequency of cooperators in the volunteer’s dilemma.

### 3.4 Experienced variability and the rest points of the replicator dynamics

The upshot of the preceding discussion in Section 3.3 is that beyond the circumstances delineated in Proposition 2 there is little hope of gaining robust insights into the effect of a change in group-size variability on the evolution of cooperation. In this section we thus focus on the impact of an increase in experienced variability for the number and location of the (stable) rest points of the replicator dynamics. We do so for games with gain sequences *d* that are either convex or concave, so that the experienced-variability effect can be signed by Proposition 1. Several case distinctions arise because convex or concave gain sequences are general enough to allow for qualitatively different dynamic regimes with zero, one, or two interior rest points. Thus, two kinds of effects may arise due to an increase or decrease in experienced variability, which we explore by considering the differences between the evolutionary dynamics for two experienced group-size distributions satisfying 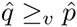. First, the number of stable rest points may stay unchanged while the location of these rests points changes. Second, the number of stable rest points might change, either via (i) a transcritical bifurcation by which an interior point collides with or emerges from the fixed point at *x* = 1, or (ii) a saddle-node bifurcation by which two interior fixed points (one stable, one unstable) are created or destroyed.

#### 3.4.1 Convex gain sequences

From Proposition 1.1 we know that condition (10) holds for convex gain sequences, so that the gain function for the more variable experienced group-size distribution 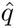 lies above the gain function for the experienced group-size distribution 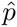. Further, by Lemma 2 the gain functions 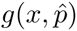 and 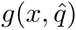 are both convex in the proportion *x* of cooperators. As a nontrivial convex function can have at most two zeros, it follows that the replicator dynamics for the two experienced group-size distributions under consideration has at most two interior rest points. Further, if the gain sequence is not only convex, but also monotonic (that is, either increasing or decreasing), so will be the expected gain function (Lemma 2), implying that in these cases there is at most one interior rest point.

If the replicator dynamics for 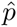 and 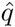 have the same number of interior rest points, then an increase in experienced variability has no effect on the stability of the rest points. For instance, if *d*_0_ < 0 holds and for both experienced group-size distributions there is a unique interior rest point, then (8) and the fact that stable and unstable rest points must alternate imply that for both 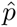 and 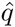 the rest point *x* = 0 is stable, the interior rest point is unstable, and the rest point at *x* = 1 is stable. Further, (10) implies that an increase in experienced variability causes the proportion of cooperators in an unstable interior rest to decrease, whereas the proportion of cooperators in a stable interior rest point increases. The left panel of Fig. 4 illustrates these assertions for the case of an increasing and convex gain sequence arising from the collective action problem in Example 2.

**Figure 4:**
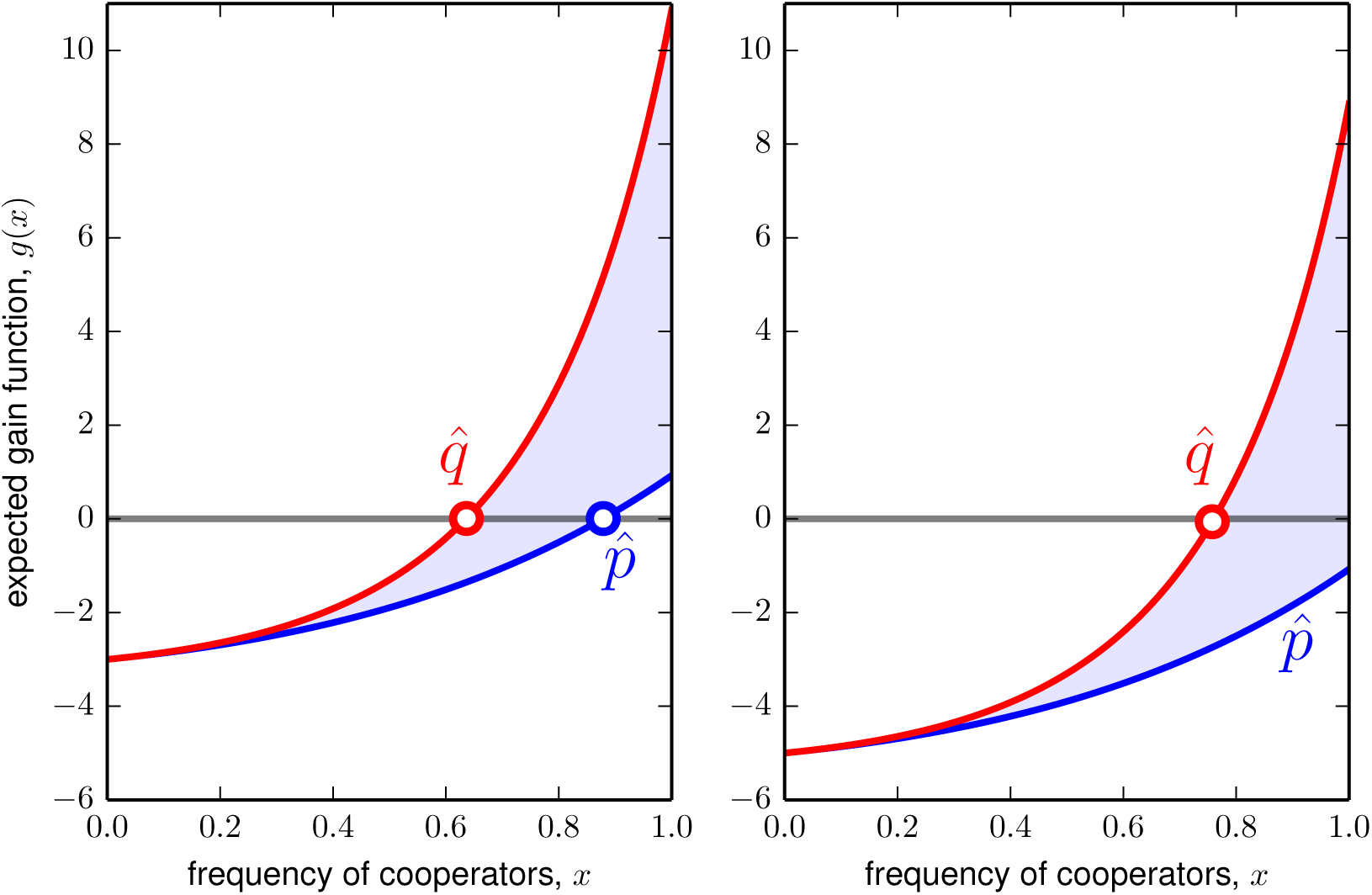
Positive experienced-variability effect for a game with increasing and convex gain sequence. The game is the public goods game with synergy or discounting of Example 2 with *u* = 1, *v* = 1.2, *w* = 1.3. The group-size distributions *p* and *q* are respectively given by a Poisson distribution Po(λ) with λ = 4 and a negative binomial distribution NB(*η*, *π*) with *η* = 1 and *π* = 2/3. With these parameters, 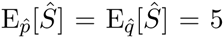 (cf. Table 1). Moreover, 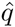 is more variable than 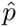 (cf. Appendix A.1). *Left. c* = 4. Increasing experienced variability from 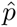 to 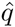 causes the unstable interior rest point (open circle) to decrease, hence increasing the basin of attraction of the fully cooperative, stable rest point *x* = 1. *Right. c* = 6. Increasing experienced variability from 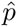 to 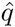 stabilizes the otherwise unstable fully cooperative rest point *x* = 1 via a transcritical bifurcation.

Depending on the sign of *d*_0_, a more variable experienced group-size distribution may either increase or decrease the number of interior rest points of the replicator dynamics for a convex gain sequence. Suppose *d*_0_ < 0 holds. We can then distinguish two cases. In the first case 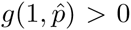 holds and the replicator dynamics for 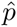 has a unique interior rest point (which is unstable). Hence (10) implies that the replicator dynamics for 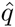 also has a unique interior rest point, so that the number of interior rest points is unchanged and the analysis from the preceding paragraph is applicable. In the second case 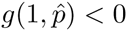 holds and there is no interior rest point. If the experienced-variability effect is sufficiently strong as to induce 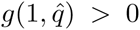, then the replicator dynamics for 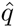 has one interior rest point and the rest point at *x* = 1 is stable, whereas the replicator dynamics for 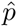 has no interior rest point and *x* = 1 is unstable. In this scenario, illustrated in the right panel of Fig. 4, the positive experienced-variability effect thus manifests itself in stabilizing a fully cooperative population via a transcritical bifurcation. In contrast, if *d*_0_ > 0 holds, then the replicator dynamics for 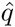 cannot have more, but might have less, rest points than the replicator dynamics for 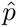. For instance, when the gain sequence *d* is convex and decreasing (as in the volunteer’s dilemma) and the inequality 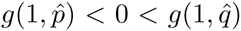 holds, then *d*_0_ > 0 implies that the replicator dynamics for 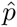 has one interior rest point, which is also the unique stable rest point, whereas the replicator dynamics for 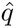 has no interior rest point and *x* = 1 is the unique stable rest point.

#### 3.4.2 Concave gain sequences

For a concave gain sequence *d* the experienced-variability effect is negative (Proposition 1.2), that is, 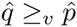 implies (11). Further, Lemma 2 shows that both 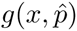 and 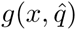 are concave in *x* and thus have at most two interior rest points.

As in the case of convex gain sequences, two scenarios are possible. First, the replicator dynamics for 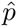 and 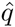 may have the same number of rest points in which case (8) implies that the stability pattern of the rest points for the two dynamics is identical and, further, (11) implies that the fraction of cooperators in an interior stable rest point is higher for the experienced group-size distribution 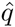, whereas the fraction of cooperators in an interior unstable rest point is higher for the experienced group-size distribution 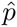. The left panel of Fig. 5 illustrates this scenario for an unimodal and concave gain function arising from the gain sequence of the collective action problem introduced in Example 2.

**Figure 5:**
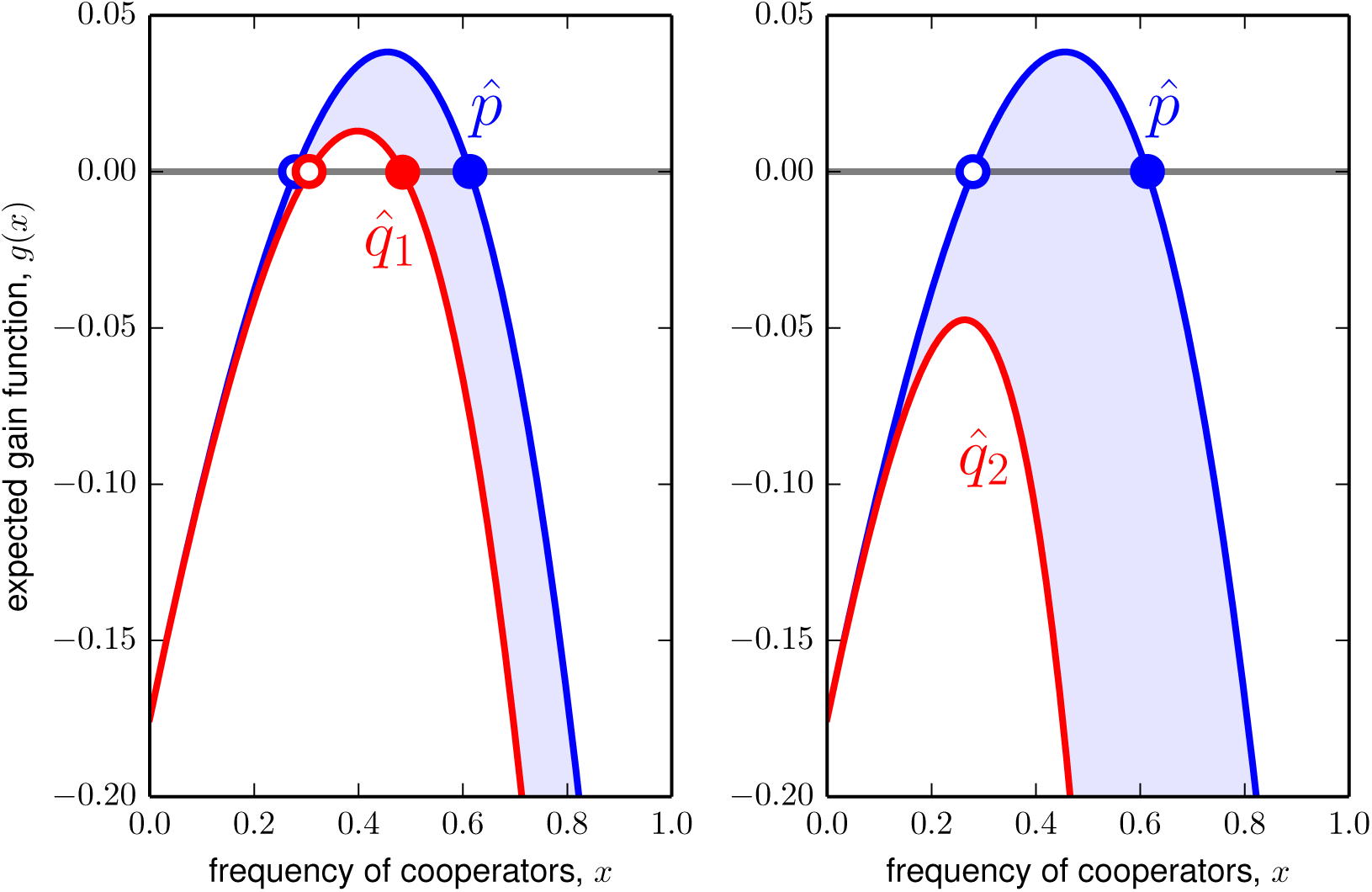
Negative experienced-variability effect for a game with unimodal and concave gain sequence. The game is the public goods game with synergy or discounting of Example 2 with *u* = 1, *v* = 1.3, *w* = 1.2, *c* = 1.175. The group-size distribution *p* is given by a Poisson distribution Po(λ) with λ = 4. The group-size distributions *q*_1_ and *q*_2_ are respectively given by negative binomial distributions ΝΒ(*η*_1_, *π*_1_) and ΝΒ(*η*_2_, *π*_2_) with *η*_1_ = 9, *η*_2_ = 1, *π*_1_ = 2/7, *π*_2_ = 2/3. With these parameters, 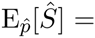 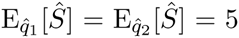 for the associated experienced group-size distributions (cf. Table 1). Moreover, 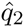 is more variable than 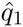 and 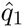 is more variable than 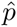 (cf. Appendix A.1). *Left.* Increasing experienced variability from 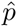 to 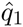 causes the unstable interior rest point (*open circle*) to increase and the stable interior rest point (*filled circle*) to decrease. The fraction of cooperators at the interior stable rest point thus decreases and its basin of attraction shrinks. *Right.* Increasing experienced variability from 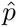 to 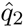 makes the gain function strictly negative. Consequently, the interior rest point disappears (through a saddle-node bifurcation) and the fully defective rest point *x* = 0 remains as the only stable rest point.

Second, the replicator dynamics for the more variable 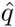 may have less (when *d*_0_ < 0) or more (when *d*_0_ > 0) interior rest points than the replicator dynamics for 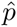. For instance, when *d*_0_ < 0 and 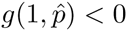 holds, then the replicator dynamics for 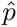 may have two interior rest points (with the first of these being unstable and the second stable), whereas with a sufficiently strong experienced-variability effect a saddle-node bifurcation occurs and the replicator dynamics for 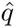 has no interior rest point. This possibility is illustrated in the right panel of Fig. 5. We note that the situation illustrated in this figure is analogous to the one considered in Bach et al. (2006), who also show that a downward shift in a unimodal concave gain function may cause the number of interior rest points of the replicator dynamics to drop from two to zero via a saddle-node bifurcation. The key difference between the scenario considered in Bach et al. (2006) and the one we consider here is that in their model the disappearance of the interior rest points is caused by a downward shift in the gain sequence *d*, whereas in our model the gain sequence is given and it is a shift in the experienced group-size distribution which induces the bifurcation causing the disappearance of the interior rest points.

## 4 Discussion

We have studied the effect of variation in group size on the evolution of cooperative behavior. Provided that variation in group size is measured according to its induced effect on the variability of the experienced group-size distribution, the model offers clear predictions: more experienced variability promotes cooperation when the payoff structure of the collective action problem implies a convex gain sequence and inhibits cooperation when the gain sequence is concave. We further showed that these variability effects can have important dynamic consequences. These include the shifting, creation, and destruction of internal equilibria and the stabilization of the full cooperative equilibrium (cf. Fig. 4 and 5). Altogether, our results add to previous work demonstrating the importance of accounting for group-size distributions in models of the evolution of social behaviors (Brännström et al., 2011; Peña, 2012).

Our analysis raises the question of which collective action problems besides the ones we have considered in Examples 1 and 2 give rise to convex or concave gain sequences. This is so for the class of club good games with accelerating or decelerating production functions considered in Peña et al. (2015). In these games defectors are excluded from the consumption of the collective good and obtain a payoff of zero. The payoff to a cooperator is *u_k_*_+1_ - *c*, where the benefit *u_j_* from obtaining the club good is increasing in the number of cooperators *j* and *c* > 0 is the cost of providing the good. Here the gain sequence is simply *d_k_* = *u_k_*_+1_ - *c*, which is convex when *u_j_* is convex (accelerating production function) and concave when *u_j_* is concave (decelerating production function). For many commonly studied collective action problems, however, the gain sequences are neither convex nor concave. Examples are (i) public goods games involving nontrivial thresholds, such that the cooperation of more than one but less than the total number of players is required to produce a collective good (Bach et al., 2006; Pacheco et al., 2009; Archetti and Scheuring, 2011), (ii) games of multiplayer reciprocity (Boyd and Richerson, 1988), and (iii) variants of the volunteer’s dilemma where the cost of providing the good is shared among cooperators (Weesie and Franzen, 1998), sometimes referred to as multiplayer snowdrift game (Zheng et al., 2007; Souza et al., 2009). No unambiguous, general conclusions concerning the effect of variable group sizes can be obtained in these cases. Instead, the gain function has to be explicitly calculated under different group size (or experienced group size) distributions in order to correctly sign the experienced-variability and the variability effects.

Throughout our analysis we have assumed that payoffs for individuals are determined by their own action (whether to cooperate or defect) and the number of cooperators in the group. All of our analysis carries over without substantial changes to the case in which the payoff consequences of own actions depend on the number of defectors (rather than the number of cooperators) in the group. Consider, for instance, the weakest-link stag hunt game (Hirshleifer, 1983). This game is like the volunteer’s dilemma, except that the cooperation of all individuals in a group is required for the benefit to be produced. To analyse this game we may consider the gains from switching as a function of the number of other individuals in the group that play defect (rather than cooperate). The resulting gain sequence is identical to the one for the volunteer’s dilemma. Consequently, Proposition 1.1 continues to apply and we may conclude that an increase (resp. decrease) in experienced variability promotes (resp. inhibits) cooperative behavior in the weakest-link stag hunt game.

As we have already noted at the end of Section 2.2, Hauert et al. (2006) assume that the benefits in their public goods game with synergy or discounting are scaled by the inverse of group-size. This implies that the gains from switching are no longer solely determined by the number of cooperators in the group but depend directly on group size. Consequently, our analysis is not directly applicable. It can be shown, however, that with such scaled benefits the very same conditions which ensure the convexity (resp. concavity) of the gain function *f* (*x*, *s*) in group size in our version of the public goods game (see Example 2) now ensure that the function *h* (*x*, *s*) = *sf* (*x*, *s*) is convex (resp. concave) in group-size. As convexity (resp. concavity) of *h*(*x*, *s*) in group size is sufficient to sign the variability effect (cf. the discussion preceding the statement of Proposition 2), we obtain the following, somewhat surprising result: with scaled benefits the variability effect can be signed under exactly the same conditions that allowed us to sign the experienced-variability effect. In particular, the results shown in Peña (2012, Fig. 2) for the scaled version of the public goods game with synergy or discounting are not limited to the particular group-size distributions considered there, but hold for arbitrary group-size distributions that are ordered by variability.

This paper has followed Peña (2012) in investigating the evolutionary consequences of variation in group size using the replicator dynamics of two-strategy multiplayer games. While this is a common approach is the literature on collective action problems (Motro, 1991; Bach et al., 2006; Peña et al., 2014), alternative approaches are possible. In particular, the very same question we are interested in has been explored by Brännström et al. (2011) in the framework of continuous strategies and adaptive dynamics (Metz et al., 1996). In contrast to us, Brännström et al. (2011) focus on a class of games in which the selection gradient (the counterpart to our gain function) is determined by the average contribution in the group, so that variability in group size has no effect on the location of the singular rest points (corresponding to the rest points of our dynamics). Rather, the effect of variation in group size in their setting reflects itself in whether evolutionary branching can occur near a singular strategy and this is the question they study. Despite such fundamental differences, the analysis in Brännström et al. (2011) shares a common feature with ours, namely that the variance of the group-size distribution is not a suitable measure of variability. The measure of variability used by Brännström et al. (2011), the average inverse group size, is consistent with our approach in the sense that more variable group-size distributions according to our definition have higher average inverse group size.

We conclude by noting that we have taken the group-size distribution to be exogenous and assumed that the experienced group-size distribution is independent of the behavior of the individuals under consideration. It would be a logical next step to extend out analysis to models in which these assumptions are relaxed. For instance, in addition to their different cooperative tendencies, individuals might vary with respect to the size of the group they would prefer to join (Powers et al., 2011) or their intrinsic ability to form groups (Garcia and De Monte, 2013). In these cases, group sizes are expected to vary endogenously in nontrivial ways. If the underlying collective action problem involves nonlinearities, the variability effects described in this paper will also arise and feed back into the evolutionary dynamics. Future work should investigate how variation in group size might affect the coevolution of group formation and cooperation in collective action dilemmas.

## Acknowledgements

This work was supported by Swiss NSF Grant PBLAP3-145860 (to J.P.).

## Appendix

### A.1 Ordering of some common (experienced) group-size distributions by their variability

Suppose the distribution *p* of *X* is negative binomial with parameters *η_x_* and *π_x_*, and the distribution *q* of *Y* is negative binomial with parameters *η_y_* and *π_y_*. Assume further that *π_x_η_x_*/(1 - *π_x_*) = *π_y_η_y_*/(1 - *π_y_*) holds, so that both random variables have the same expected value (cf. Table 1). Whitt (1985) demonstrates that the relationship *q* ≥*_v_ p* is then implied if the inequality

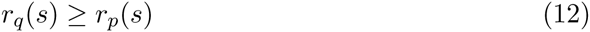

holds for all *s* ≥ 1, where *r_p_*(*s*) and *r_q_*(*s*) denote the indices of relative log-concavity of the two distributions, which can be calculated as (see Whitt, 1985, Example 6)

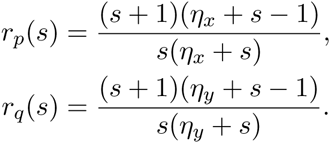

A straightforward calculation shows that condition (12) is satisfied for all *s* ≥ 1 when *η_x_* ≥ *η_y_* holds. Consequently, we have *q* ≥*_v_ p* if *η_x_* ≥ *η_y_* is satisfied. Further, as the Poisson distribution can be obtained as the limit of the negative binomial distribution for *η* → ∞ and the geometric distribution corresponds to the negative binomial with *η* = 1, it follows that (provided the expected values are identical) a geometric distribution is more variable than any negative binomial distribution with *η* > 1 and every negative binomial distribution is more variable than a Poisson distribution. In particular, any two of the experienced group-size distributions appearing in Table 1 are ordered by variability when they have the same expected value.

These kinds of comparisons can be extended to many other familiar distributions. For instance, the Waring distribution considered in Peña (2012) as a group-size distribution is a mixture of geometric distributions (Johnson et al., 2005, p. 290) and is thus (Whitt, 1985, Example 6) more variable than the geometric distribution with the same expected value as the Waring distribution under consideration. Similarly, it is immediate from Johnson et al. (2005, Eq. 7.21, p. 307) that the logarithmic distribution featured in Table 1 is more variable than the geometric distribution.

### A.2 Proofs

#### Proof of Lemma 1

The polynomial in Bernstein form of degree *n* of the sequence *d* = (*d*_0_, *d*_1_, *d*_2_,...) is

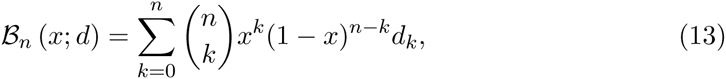

so that from (3) we have

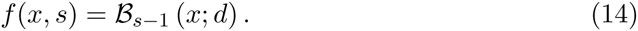

By (14) we have

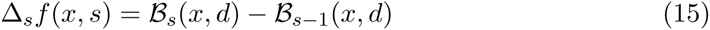

and

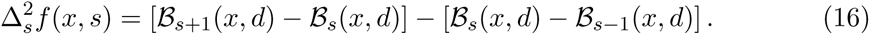

Motro (1991) shows (cf. the proof of part (ii) of the proposition in his appendix) that

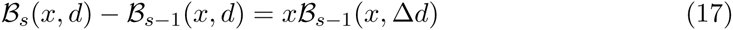

holds for *x* ∈ [0, 1], *s* ≥ 1 and all sequences *d*. Observing that the polynomial in Bernstein form appearing on the right side of (17) is positive (negative) when all its coefficients are positive (negative) it follows from (15) that for increasing (decreasing) *d* we have that Δ*_s_f* (*x*, *s*) is positive (negative) for all *x* ∈ [0, 1] and *s* ≥ 1. This establishes that *f* (*x*, *s*) is increasing (decreasing) in *s* when *d* is increasing (decreasing).

Applying (17) to both terms in square brackets in (16) and simplifying we obtain

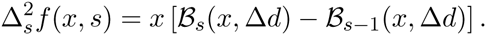

As (17) holds for all sequences *d*, we can apply it with Δ*d* in place of *d*, to obtain

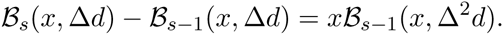

Combining the previous two equalities yields

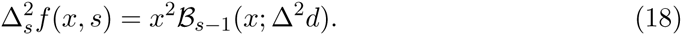

As the right side of (18) is positive (negative) if *d* is convex (concave), this establishes that *f* (*x*, *s*) is convex (concave) in *s* if *d* is convex (concave).

#### Proof of Proposition 1

Let 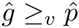 and let *d* be convex. From Lemma 1, convexity of *d* implies that *f* (*x*, *s*) is convex in *s*. Because 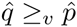 implies that the inequality 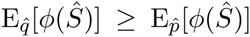 holds for all convex functions *φ*, it follows that 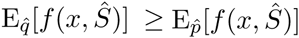 holds for all *x* ∈ [0,1]. Substituting the definition of the gain function (4) into this inequality, we obtain (10). Consequently, the experienced variability effect is positive when *d* is convex.

When *d* is concave, Lemma 1 implies that *f* (*x*, *s*) is concave in *s*, so that -*f* (*x*, *s*) is convex in *s*. Hence 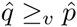 yields that the inequality 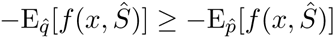 holds for all *x* ∈ [0, 1]. Multiplying both sides of the inequality by −1 and using the definition of the gain function (4), we obtain (11). Consequently, the experienced variability effect is negative when *d* is concave.

#### Proof of Proposition 2

We show that for increasing and convex *d*, *q* ≥*_v_ p* implies (10), thus establishing the first part of the proposition. (As in the proof of Proposition 1, the result for the case in which *d* is decreasing and concave follows by an analogous argument.)

Using (6), we may write

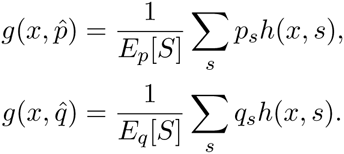

As *q* ≥*_v_ p* implies *E_p_*[*S*] = *E_q_* [*S*], it follows that the variability effect is positive if

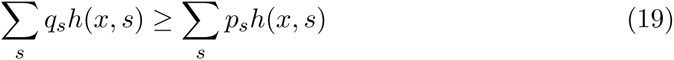

holds for all *x*. By definition of the convex order, (19) is satisfied whenever *h*(*x*, *s*) is convex in *s*, so that it suffices to establish this property.

If *d* is increasing and convex, then *f* (*x*, *s*) is increasing and convex in group-size *s* (Lemma 1). Using the definition *h*(*x*, *s*) = *sf* (*x*, *s*), a straightforward calculation shows that

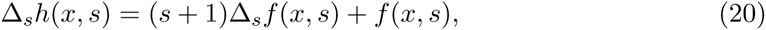

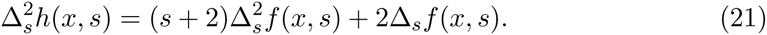

Because *f* (*x*, *s*) is increasing and convex in group size, it satisfies Δ*_s_ f* (*x*, *s*) ≥ 0 and 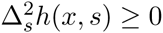, so that (21) implies 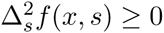. Hence, *h*(*x*, *s*) is convex.

## Footnotes

1 Here and throughout our formal analysis we focus on the effects of an increase in (experienced) variability as the corresponding results for the effects of a decrease in (experienced) variability are easily inferred as they are simply opposite in sign. For instance, Proposition 1 can be read as the statement that less variation in experienced group size inhibits cooperation when the gain sequence is convex and promotes cooperation when the gain sequence is concave.

2 We refrain from making stronger assumptions on the support of the group-size distribution —such as imposing a lower and/or upper bound— to accommodate commonly considered models for group-size distributions that we use for illustrative purposes.

3 The gain sequence for the volunteer’s dilemma is the limit case of the gain sequence in (2) for *v* = *w* → 0.

4 It is not difficult, but tedious, to show that the expected gain function is determined by the first two moments of the experienced group-size distribution if and only if the gain sequence takes the form *d_k_* = *α* + *βk* + *γk*^2^ for some parameters *α*, *β*, and *γ*. In the context of a public goods game with constant cost *c* > 0 of contributing to the public good, the gain sequence will take this form if and only if the benefit of the public good is a polynomial of degree no larger than 3 in the number of contributors. For the expected gain function to be determined by the first two moments of the group-size distribution the additional restriction *γ* = 0 is required.

